## Supplemental Figures and Legends for "Defining Cellular Diversity at the Swine Maternal-Fetal Interface Using Spatial Transcriptomics and Organoids"

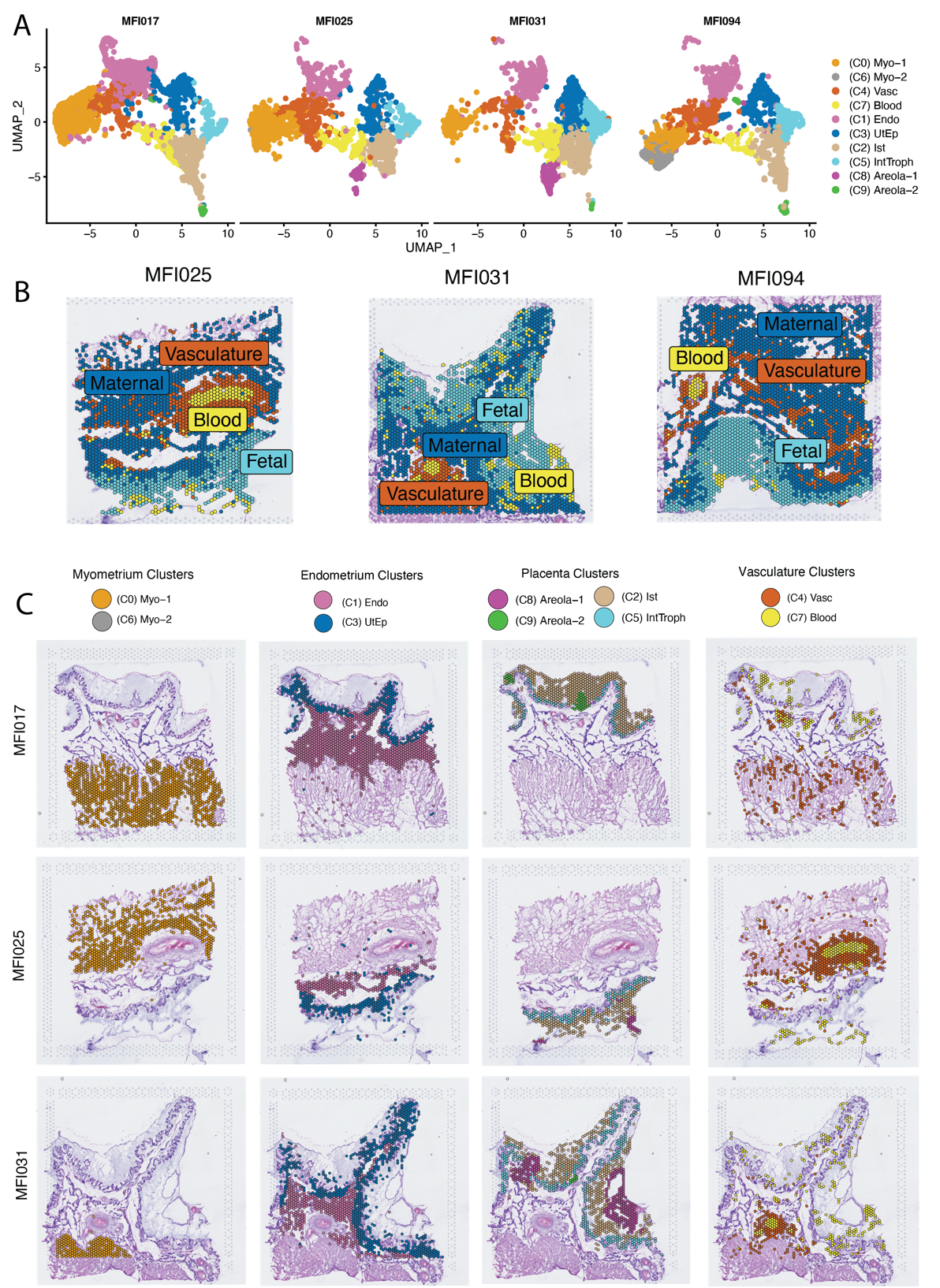

**Supplemental Figure 1:** **A)** UMAP split by sample (MFI). Dots indicate individual visium spots. **B)** Spatial Dimplot showing separation of maternal and fetal components. Representative maternal-fetal Interface shown. C) Spatial DimPlot separated by sample type and histologic structures showing the localization of UMAP cluster populations.

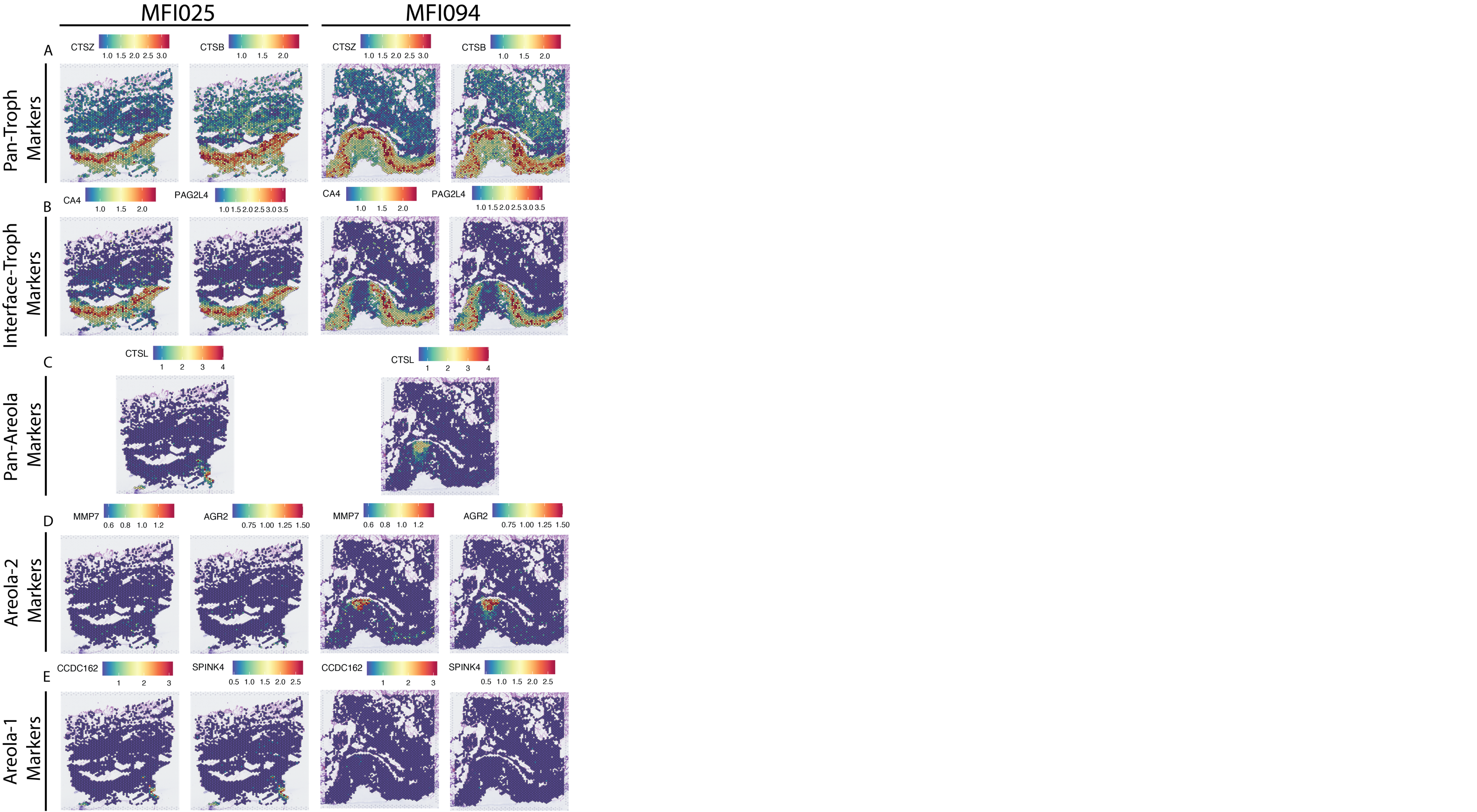

**Supplemental Figure 2:** **A-E)** Spatial Feature plots of known and novel markers of various trophoblast populations, split by sample (MFI). Various populations include A) Pan trophoblast markers, .) Interface trophoblast markers, C) Pan-Areola markers, D) Areola-2 markers, E) Areola-1 markers.

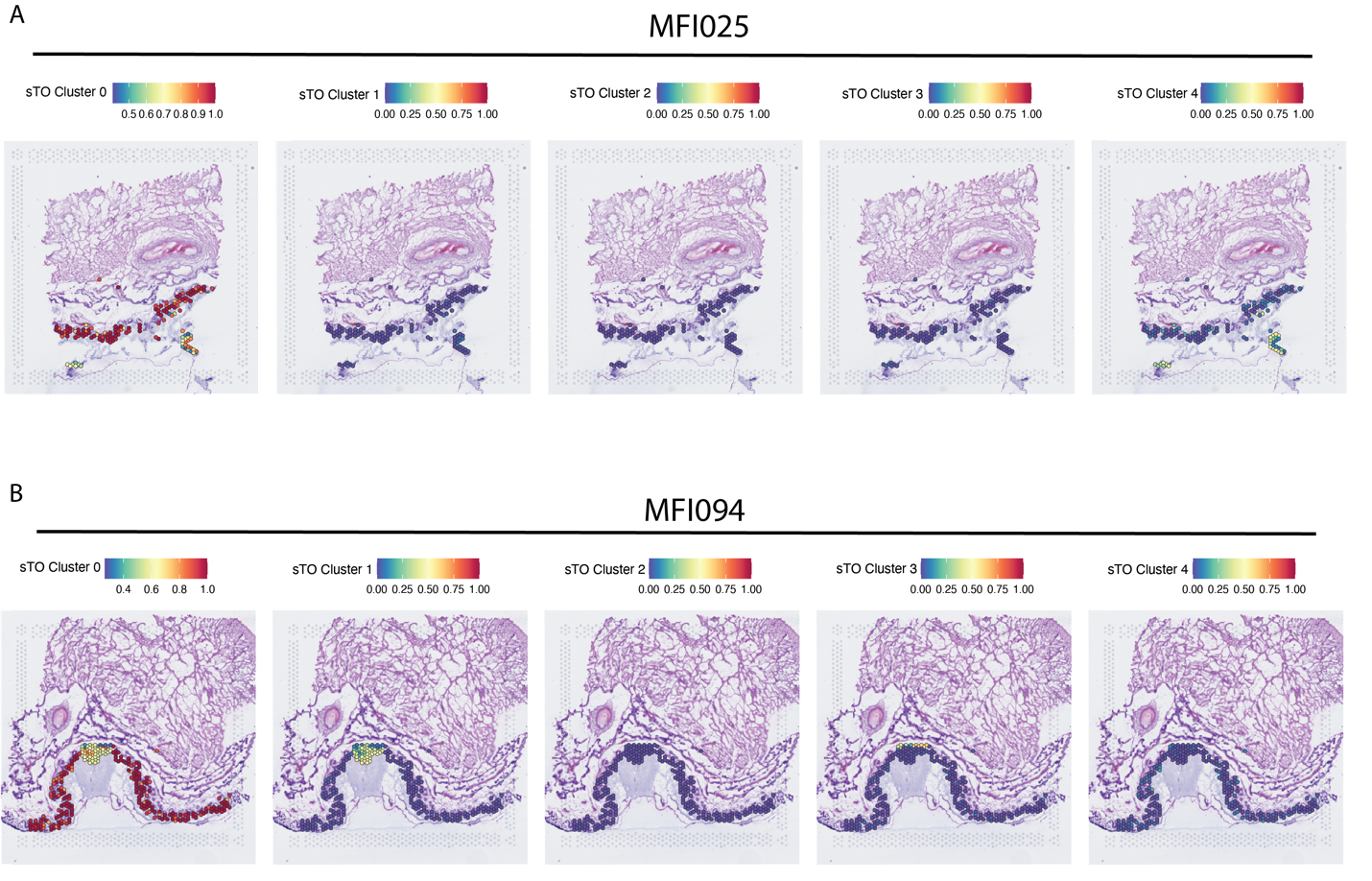

**Supplemental Figure 3: A-B)** Spatial feature plot showing prediction scores for each spot in the spatial dataset subsetted by sTO cluster. Color shown indicates probability that the classified sTO cluster is localized to a given position within the spatial dataset. A) Scoring for MFI025. B) Scoring for MFI094.

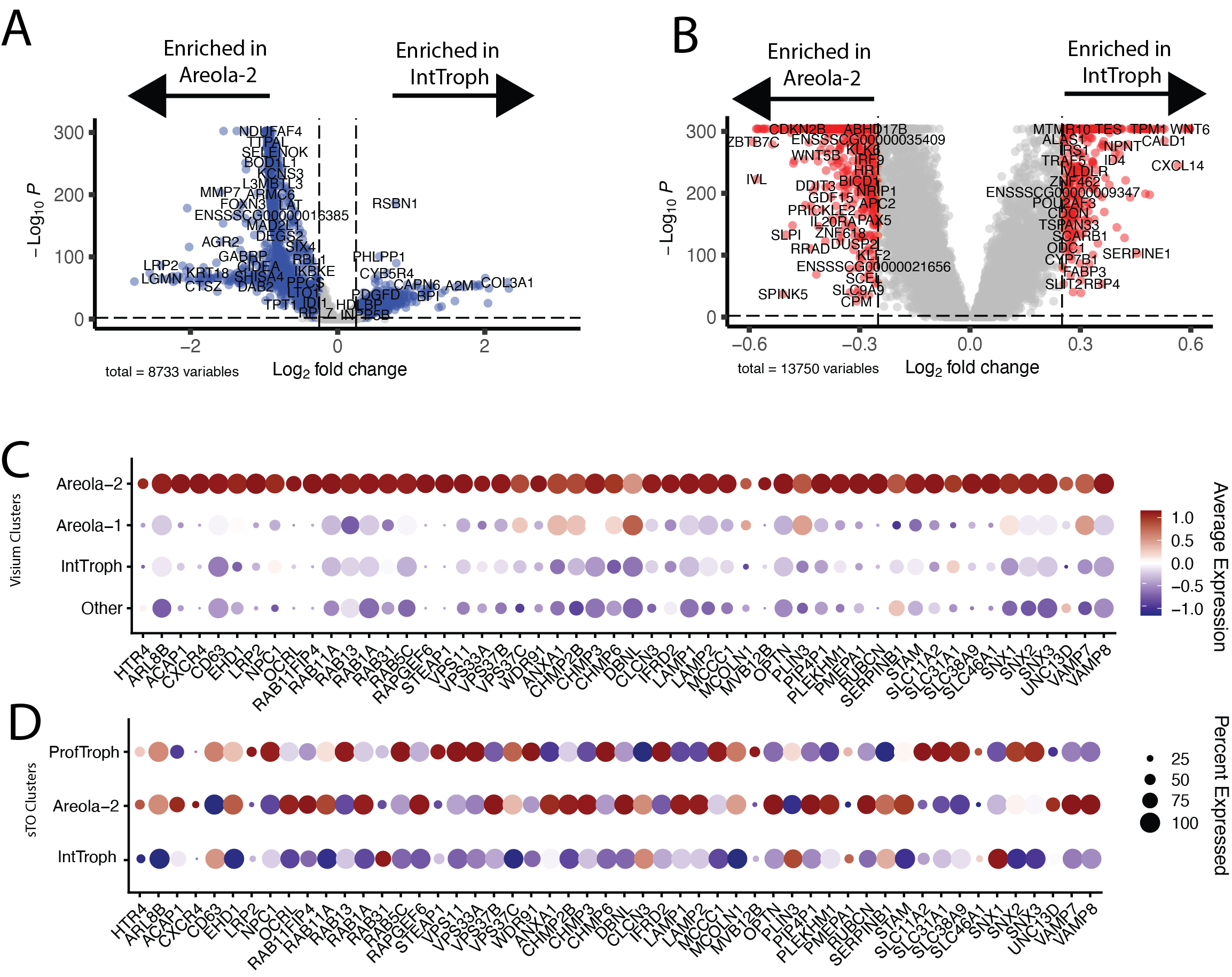

**Supplemental Figure 4: Differential Expression Highlights the Distinct Functional Niches of Interface and Areola-2 Trophoblasts.** **A)** Volcano plot showing differentially expressed genes in interface-trophoblasts and areola-2 trophoblasts within the Visium Spatial Transcriptomics data. Significant differences (p<0.01, Fold-change > 0.25) are shown in blue. **B)** Volcano plot showing differentially expressed genes in interface-trophoblasts and areola-2 trophoblasts within the sTO single cell transcriptomics data. Significant differences (p<0.01, Fold-change > 0.25) are shown in red. **C-D)** Dotplots showing the expression of endosome specific genes in both Visium (C) and sTO datasets (D). Plots showing Interface-trophoblast enriched GO-Terms as calculated by DAVID pathway analysis (significant difference = p<0.05) for Visium spatial transcriptomics (C) and sTO single cell (D). Color of dots denote average expression, whereas size represents percentage of cells in a cluster expressing the gene of interest.

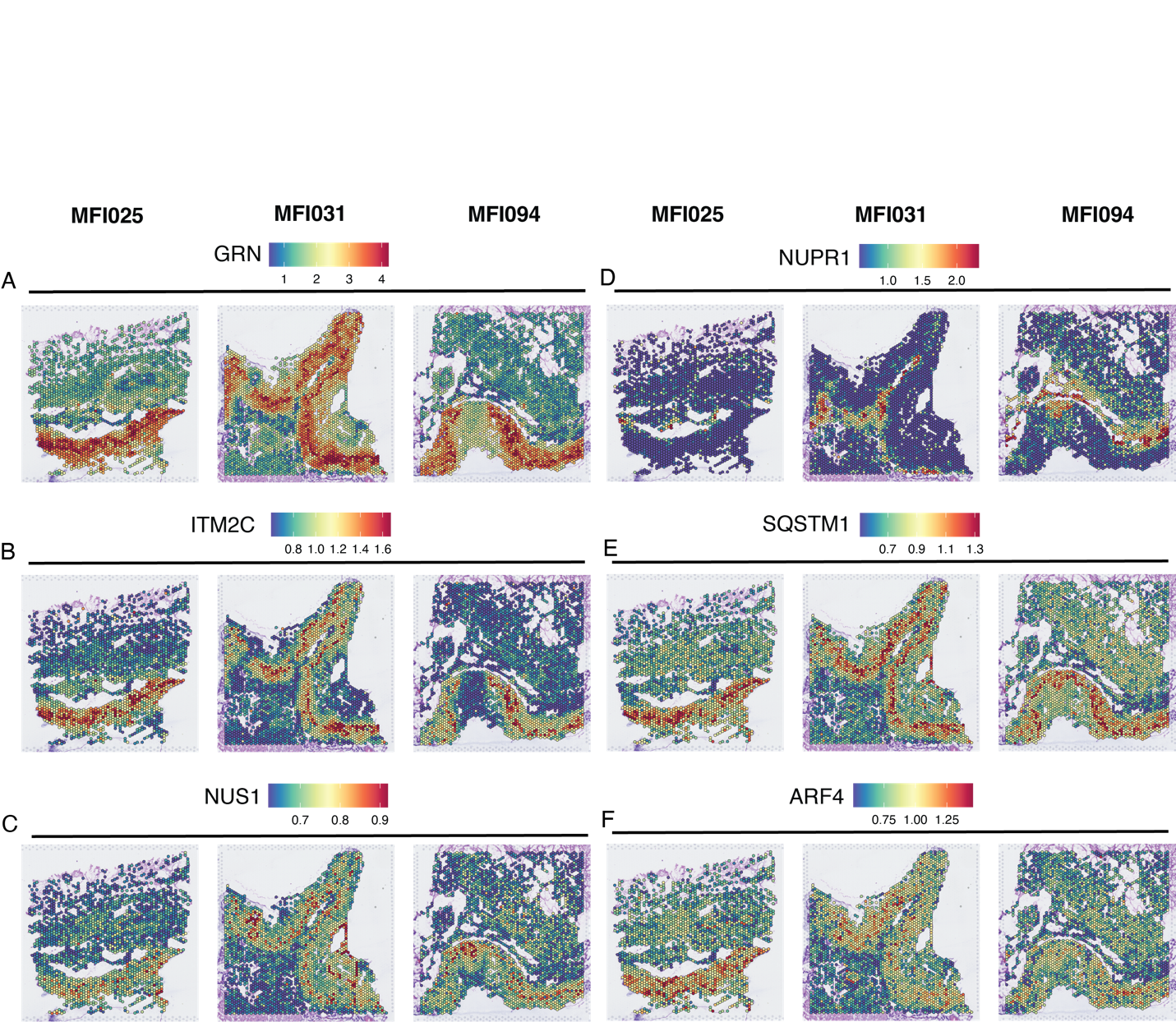

red being high and blue being low contribution level.

**Supplemental Figure 5: A-F)** Spatial FeaturePlots showing genes that are enriched during the transition to Areola-2 trophoblasts and their expression pattern within our spatial transcriptomics dataset, split my sample (MFI). A.) GRN, B.) ITM2C, C.) NUS1, D.) NUPR1, E.) SQSTM1, and F.) ARF4.
